# Inferring extrinsic factor-dependent single-cell transcriptome dynamics using a deep generative model

**DOI:** 10.1101/2024.04.01.587302

**Authors:** Yasuhiro Kojima, Yuko Arioka, Haruka Hirose, Shuto Hayashi, Yusuke Mizuno, Keiki Nagaharu, Hiroki Okumura, Masato Ishikawa, Kohshi Ohishi, Yutaka Suzuki, Norio Ozaki, Teppei Shimamura

**Author notes:** Equal contribution (Yasuhiro Kojima, Yuko Arioka).

## Abstract

RNA velocity estimation helps elucidate temporal changes in the single-cell transcriptome. However, current methodologies for inferring single-cell transcriptome dynamics ignore extrinsic factors, such as experimental conditions and neighboring cell. Here, we propose ExDyn—a deep generative model integrated with splicing kinetics for estimating cell state dynamics dependent on extrinsic factors. ExDyn enables the counterfactual inference of cell state dynamics under different conditions. Among the extrinsic factors, ExDyn can extract key features which have large effects on cell state dynamics. ExDyn correctly estimated the difference in dynamics between two conditions and showed better accuracy over existing RNA velocity methods. ExDyn were utilized for unveiling the effect of PERK-knockout on neurosphere differentiation, hematopoietic stem cell differentiation driven by chromatin activity and the dynamics of squamous cell carcinoma cells dependent on colocalized neighboring cells. These results demonstrated that ExDyn is useful for analyzing key features in the dynamic generation of heterogeneous cell populations.

## 2 Introduction

Cells are dynamic systems that respond to both intrinsic and extrinsic factors by altering their internal state. Single-cell transcriptome analysis has revealed the heterogeneity of internal cell states in various biological processes. A comparative study of multiple experimental conditions reported the enrichment of certain cell states under specific conditions. Recent technological advancements in single-cell transcriptomics have enabled the incorporation of batched experimental conditions and single-cell wise information, such as protein expression [45], chromatin accessibility [36], and spatial information [7, 41]. These advances also allow for the analysis of single-cell transcriptome augmentations with extrinsic and intrinsic factors. However, the effects of these factors on cell state dynamics and heterogeneity remain largely unclear because single-cell transcriptomics is invasive and provides only a snapshot of cell states.

La Manno et al. developed RNA velocity—a method that utilizes splicing kinetics to reconstruct temporal changes in gene expression—which has enabled the data-driven analysis of transition relationships within heterogeneous populations [28]. Many methodologies were proposed to improve the accuracy and applicability of RNA velocity estimation [3,11,17,29]. Several of them adopted the framework of a deep generative model [9,19], which is a key technology for single cell omics analysis. However, no methodology has been proposed to uncover the regulation of cell state dynamics by extrinsic factors, such as experimental conditions and signaling from neighboring cells.

To reveal the regulatory relationship between extrinsic factors and cell state dynamics, we propose ExDyn—a method for inferring extrinsic factor-dependent cell state dynamics. Using a deep generative model of the unspliced transcriptome based on latent cell state dynamics and the splicing kinetics equation, we conducted variational inference of the dynamics from the extrinsic factors and the latent cell state using a neural network. In other words, the unspliced transcriptome that reflects the near future of the spliced transcriptome was estimated from the current cell states and extrinsic factors under the constraint of splicing kinetics. By leveraging the optimized cell state dynamics associated with the extrinsic factors, ExDyn 1) provides a counterfactual estimate of cell state dynamics under different conditions for an identical cell state, 2) identifies the bifurcation point between experimental conditions, and 3) performs a principal mode analysis of the perturbation of cell state dynamics by multivariate extrinsic factors, such as epigenetic states and cellular colocalization. Using the corresponding spatial transcriptome observations, we analyzed the ability of ExDyn to identify candidate cell–cell interaction that govern cell state transitions. We investigated the ability of ExDyn to estimate differential cell state dynamics using simulated data and demonstrated its performance superior to other RNA velocity tools. We examined the impact of Protein kinase R-like endoplasmic reticulum (PERK) knockout (KO) on the cell state dynamics of a neurosphere population. Finally, we explored the roles of epigenetic states and neighboring cells on cell state dynamics in hematopoiesis and squamous cell carcinoma. Our findings indicated that ExDyn provides a promising tool for exploring the regulation of cell state dynamics by various extrinsic factors.

## 3 Results

### 3.1 Overview of ExDyn

ExDyn is a deep generative model that provides extrinsic factor-dependent cell state dynamics from spliced and unspliced transcriptome observations (**Fig. 1**). While the spliced transcriptome is derived by nonlinear transformation using a neural network, the unspliced transcriptome is assumed to be dependent on cell state dynamics through a constraint of RNA velocity. Using variational inference, we approximated the posterior distribution of cell state dynamics using a neural network that takes internal cell states and extrinsic factors as inputs. Briefly, the variational posterior distribution of cell state dynamics was optimized based on the splicing kinetics equation to maximize consistency with the unspliced transcriptome. We noted that using a conditional variational autoen-coder (cVAE) [26], such as in an existing deep generative model [32], removed the batch effects from the cell state space so that the cell state dynamics were defined on a consistent latent space across experimental batches. The optimized variational posterior distribution retained information about the complex relationships between cell state dynamics and extrinsic factors utilized for downstream analysis. The estimated relationships between the cell state dynamics and discrete experimental conditions allowed for a single-cell-wise comparison of gene expression dynamics between experimental conditions, which we used to identify the drivers of the dynamic differences and bifurcation points between condition-specific cell populations. For more complex factors, such as neighboring cells and chromatin accessibility, the principal mode analysis of cell state dynamics perturbation by extrinsic factors was conducted through singular value decomposition (SVD) of the Jacobian matrix, which was derived by the differentiation of cell state dynamics by extrinsic factors. This analysis provided the principal dynamic changes induced by the variation in the extrinsic factors and the key features of the extrinsic factors contributing to the principal patterns, which we used to explore the cellular heterogeneity associated with the extrinsic factors. From the corresponding spatial transcriptome data, we further explored cell–cell interactions and signaling molecules that drive the dynamics toward a specified cell population.

**Figure 1:**
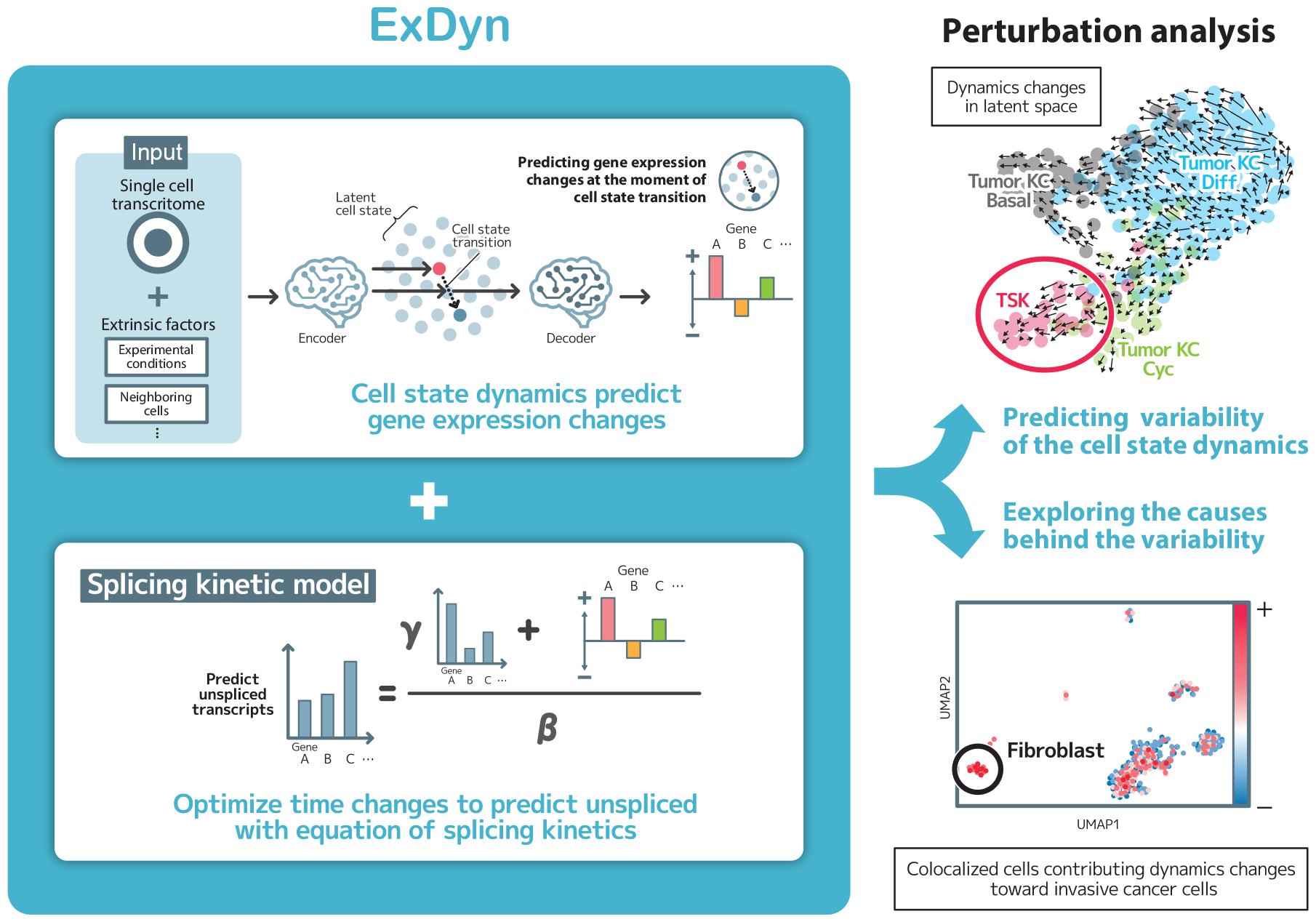
Overview of the proposed method. We assumed stochastic dynamics of VAE cell states dependent on state and condition. The dynamics were optimized based on the splicing kinetics model to be consistent with the observed unspliced transcriptome. Using the optimized model, we can assess how the cell state dynamics is perturbed by the extrinsic factors and which feature of the extrinsic factors contribute to the perturbations.

### 3.2 ExDyn accurately estimates the differential dynamics between conditions

We validated the performance of ExDyn using simulated single-cell RNA sequencing (scRNA-seq) data under two different conditions. The source cell population (cell type 0) was differentiated into cell type 1 under condition 1 and cell type 2 under condition 2 by varying the transcription rate of the master regulators. The transcription rate of transcription factor 9 was higher in condition 1, whereas that of transcription factor 18 was higher in condition 2. We applied ExDyn to estimate the state dynamics of each cell under each condition and found that the estimated dynamics were directed from the source population to both target populations under the conditions where the cells were originally observed (**Fig. 2-A**). We evaluated the dynamics in both conditions for all cells and found that most cells with common states between conditions 1 and 2 differentiated into cluster 1 under condition 1 (**Fig. 2-B**), while they differentiated into cluster 2 under condition 2 (**Fig. 2-C**). The estimated expression patterns and velocity under the two conditions confirmed that the positive velocity of genes specifically expressed in cell type 1 was associated with intermediate cell states between the source population and cell type 1 only under condition 1, while this was observed for cell type 2 under condition 2 (**Fig. 1-A,B,C in Supplementary**) . Therefore, ExDyn can not only recover the expected cell state dynamics in the conditions where the cells were observed but also predict the dynamics under varying conditions. We analyzed cells with large differences in transcriptome dynamics between conditions. The norm of the differential RNA velocity across genes for each cell showed that the dynamic difference was larger at the bifurcation point than in the root or terminal cell populations (**Fig. 2-D**). Averaged cell state dynamics in each condition across cells with the top 20% conditional difference in gene expression velocity were directed to terminal states in corresponding conditions (**Fig. 2-E**), suggesting that the differential cell state dynamics were correctly estimated at the bifurcation point between condition-specific populations. To identify genes with large conditional differences of velocity, we averaged the difference in gene expression velocity between the conditions across cells with the top 10% conditional differences in gene expression velocity. We found that, among 100 genes, the average dynamic differences of master regulators 9 and 18 were the 4th and 4th largest under conditions 1 and 2, respectively (**Fig. 2-F**). These results show that the bifurcation point effectively represents the dynamic difference between the conditions and can aid in identifying the molecular machinery of bifurcation.

**Figure 2:**
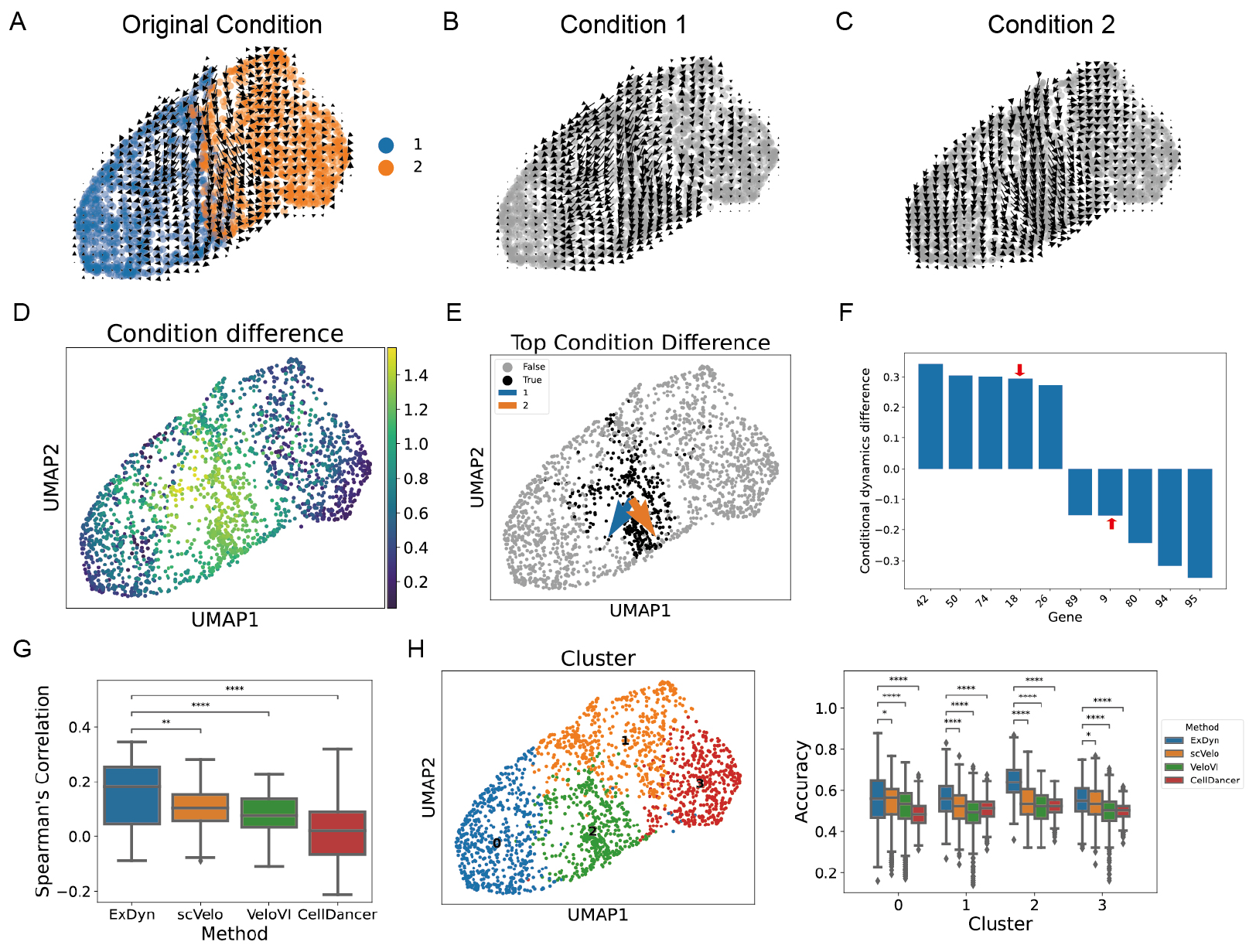
Differential cell sate dynamics in simulated dataset. **A, B, C**, Projection of cell state dynamics into UMAP embeddings of latent cell states in the conditions where the cells were originally observed (**A**), condition 1 (**B**), and condition 2 (**C**). **D**, L2-norm of the difference in gene expression velocity between the conditions. **E**, Averaged cell state dynamics in condition 1 and 2 (blue and orange arrows) of cells with top 20% gene expression velocity changes (black dots) between conditions. **F**, Conditional dynamic differences in genes quantified based on the average gene expression velocity difference between conditions across cells represented by black dots in **E**. Red arrow indicates master regulators (gene 9 and gene 18 in condition 1 and 2 respectively). **G**, Gene-wise Spearman’s rank correlation coefficients between the estimated RNA velocity and ground-truth RNA velocity across cells. We tested whether the correlation coefficients varied significantly between methods using a Mann–Whitney U test and corrected the p-values using the Benjamini–Hochberg method. Ns: 0.05 *< p ≤* 1, *: 0.01 *< p ≤* 0.05, **: 0.001 *< p ≤* 0.01, ***: 0.0001 *< p ≤* 0.001, ****: *p ≤* 0.0001. **H**, Cluster-wise distribution of cell-wise accuracy of RNA velocity estimated using ExDyn and other RNA velocity methods (scVelo, VeloVI, and CellDancer). The left panel represents the clusters in UMAP embeddings of latent cell states, and the right panel represents the distribution of cell-wise accuracies for each cluster. We tested whether the accuracy was significantly different between ExDyn and the other methods in the same manner as in **G**.

Using the ground-truth RNA velocity data, we evaluated and compared the accuracy of the estimated RNA velocity with that of other methods (scVelo [3], VeloVI [19], and CellDancer [29]). A gene-wise Spearman’s rank correlation between the estimated and ground-truth RNA velocities indicated that ExDyn had the highest accuracy among the methods (**Fig. 2-G**). As a cell-wise accuracy metric, we calculated the proportion of genes that had consistent signs in the estimated and ground-truth RNA velocities for each cell. The superiority of ExDyn compared to the other methods was most distinct at the bifurcation between the conditions, whereas the accuracy of ExDyn was also significantly higher than that of the other methods in other cell populations (**Fig. 2-H**), (**Fig. 1-D in Supplementary**) . This was presumably because the cell state dynamics at the bifurcation point were distinctively different between the conditions even if the cell states itself are quite similar, for which the other methods are expected to estimate similar transcriptome dynamics. These results demonstrated that ExDyn achieves a higher accuracy in RNA velocity estimation than existing methods by accounting for the differences in gene expression dynamics between conditions.

### 3.3 ExDyn captures the effect of PERK KO on cell state dynamics in neurospheres

We applied ExDyn to the scRNA-seq data of iPS cell-derived neurospheres from PERK-KO and WT mice. PERK is essential for the adaptation of cells to ER stress [22]. We found that the latent cell state learned by ExDyn distinguished astrocyte-like, neuron-like, and oligodendrocyte-like populations from stem cell populations (**Fig. 3-A**), (**Fig. 2-A in Supplementary**) . Within the stem cell population, we observed distinct cell cycle phases: G2M, G1, and S phases (**Fig. 3-A**), (**Fig. 2-B in Supplementary**) . Clustering analysis also distinguished the subpopulations of astrocyte-like (Astrocyte-like 2) and S phase cells (S phase 2) (**Fig. 2-C in Supplementary**), which were enriched in the KO and WT groups, respectively (**Fig. 3-B**). Gene ontology enrichment analysis of the differentially expressed genes revealed that hypoxia-related genes were upregulated in the Astrocyte-like 2 population (**Fig. 3 in Supplementary**) . The estimated dynamics under the conditions where the cells were originally observed recapitulated not only the differentiation toward several cell lineages, such as astrocytes, neurons, and oligodendrocytes, but also the cycling of the stem cell population (G2M phase 1 *→* S phase 1, S phase 2 *→* G2M phase 2) (**Fig. 3-C**). On the other hand, the dynamic difference between the conditions showed that the cell state dynamics were biased toward the Astrocyte-like 2 and S phase 1 population in the PERK-KO and WT conditions, respectively (**Fig. 3-D**). Furthermore, the Astrocyte-like 2 and S phase 1 cell populations exhibited a large differences of the cell state dynamics (**Fig. 3-E**). The estimated transition between the clusters showed that the flux from the G2M phase to the WT-enriched S phase and the abundance of the astrocyte-like population were largely decreased in the PERK-KO group compared to those in the WT group (**Fig. 3-F**). Therefore, ExDyn successfully estimated the dynamic difference between PERK-KO and WT neurospheres, which elucidated the changes in cell population abundance between conditions.

**Figure 3:**
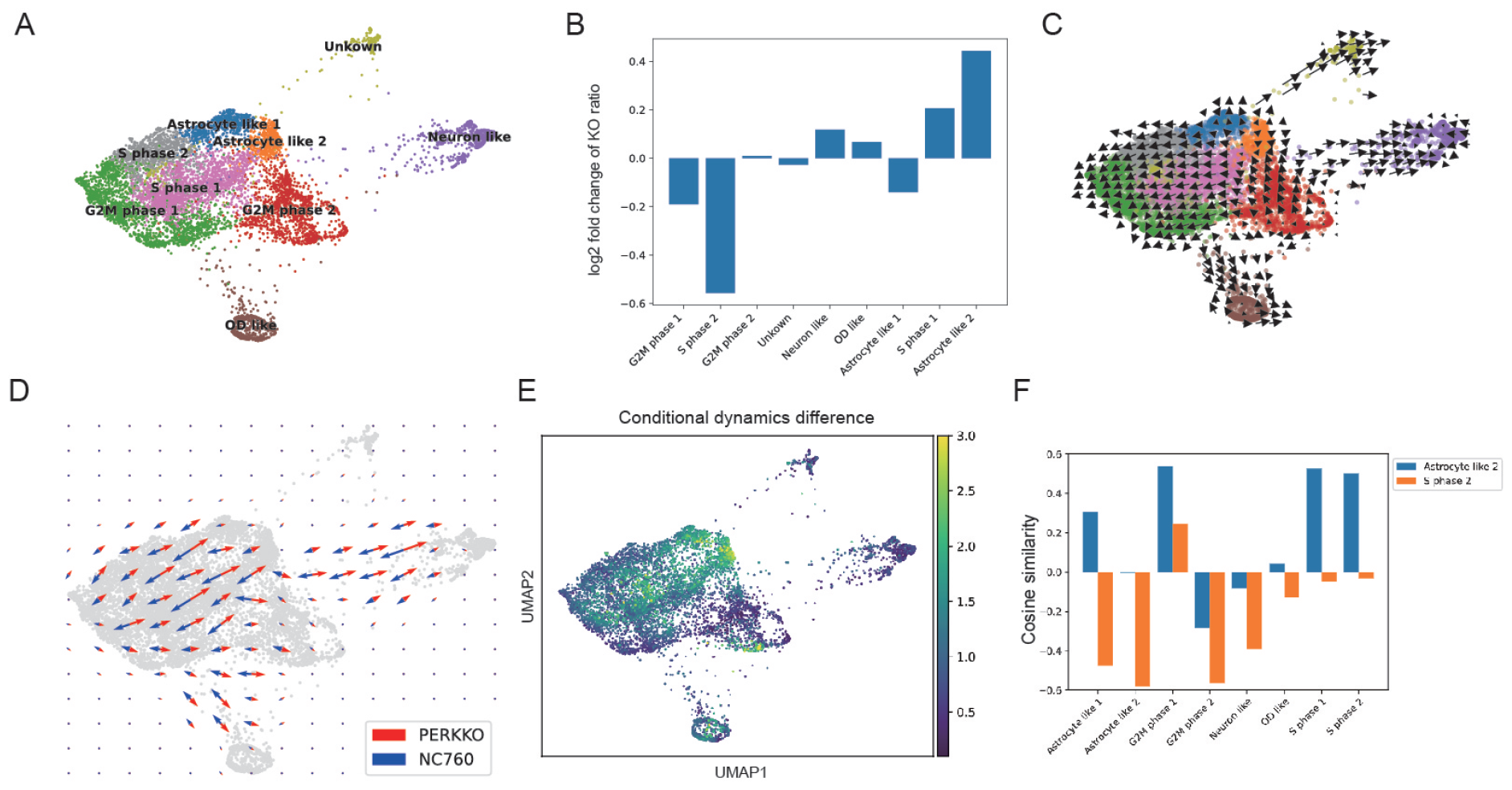
Cell state dynamics difference between PERK KO and WT in neurospheres. **A**, UMAP embeddings of latent cell states estimated by ExDyn. The cells are colored according to the annotations determined using the signature scores in (**Fig. 2-A,B in Supplementary**) . **B**, Log2 fold changes in the proportion of PERK-KO cells between each cluster and the total population. **C**, Cell state dynamics projected into two-dimensional UMAP embeddings of latent cell states in original condition. **D**, Differences in cell state dynamics between PERK-KO and WT conditions projected into two-dimensional UMAP embeddings of latent cell states. **E**, Magnitude of cell state dynamic differences between PERK-KO and WT groups for each cell. The magnitude is quantified by the L2-norm of the difference in transcriptome velocity between the conditions. **F**, Cosine similarity between relative positions of mean cell states for clusters and cell state dynamic differences between PERK-KO and WT groups. We evaluated the positions from all the clusters relative to Astrocyte-like 2 (enriched in PERK KO) and S phase 2 (enriched in WT) clusters, while the cell state dynamic differences between the conditions were calculated for all the clusters.

### 3.4 Identification of regulator for PERK KO specific astrocyte-like population

We explored the molecular mechanisms associated with the generation of the PERK KO-specific population by identifying the branching point and candidate regulators (genes with large conditional differences at the branching point) that contribute to cell differentiation. First, we extracted cells with conditional differences in gene expression dynamics (between WT and KO mice) larger than the 90% quantile and conducted a clustering analysis (**Fig. 4-A**). Directional analysis of conditional difference in cell state dynamics revealed that the cell state dynamics in cluster 3 was strongly directed toward KO-specific astrocytes in the KO condition (**Fig. 4-B**). We evaluated the conditional differences in each gene of cluster 3 and found that the positively regulated genes were strongly expressed in the KO-specific astrocyte population, while the negatively regulated genes were weakly expressed (**Fig. 4-C**). Among the upregulated genes, MIAT was upregulated under ER stress in a breast cancer cell line [51] and PFKB4 participated in the metabolic response of cancer cells to hypoxia [10]. We experimentally validated whether PFKB4 participated in the up-regulation of hypoxia-related genes in the KO-specific Astrocyte-like 2 population. We confirmed that the expression of the hypoxia-related genes BNIP3 and PDK1 was upregulated in PERK-KO mice (**Fig. 4-D**) and downregulated by siRNA knockdown of PFKB4 (**Fig. 4-E, F**). These results suggest that PFKB4 upregulates the expression of hypoxia-related genes in the KO-specific Astrocyte-like 2 population and demonstrate the ability of ExDyn for identifying the molecular mechanisms underlying condition-specific cell differentiation.

**Figure 4:**
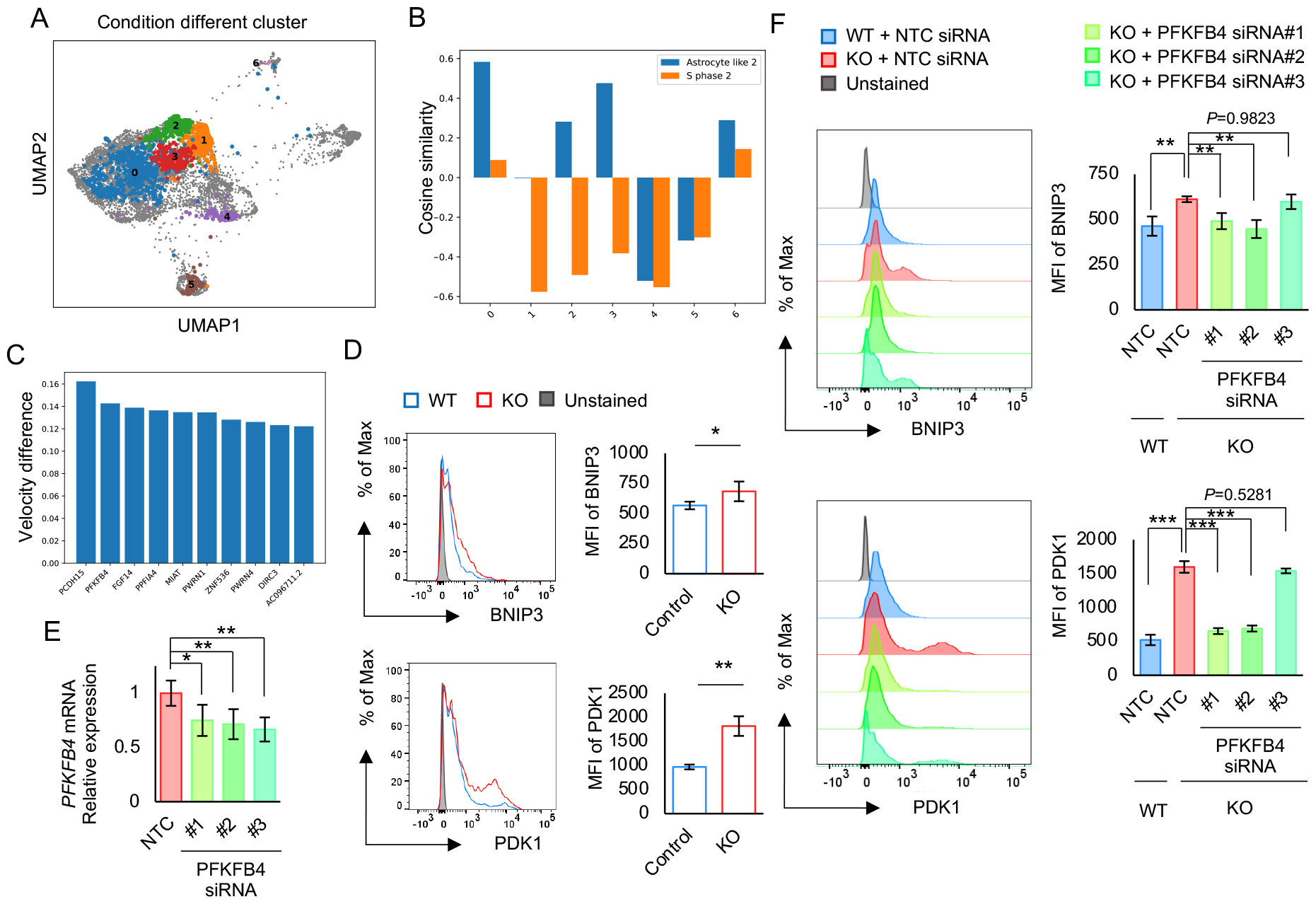
Molecular mechanism for KO-enriched astrocyte-like populations. **A**, Clusters of cells with large conditional dynamic differences (top 10%). The other cells are represented by gray dots. **B**, Cosine similarity between relative positions of cell states and cell state dynamic differences between PERK-KO and WT groups. We evaluated the relative positions from all the clusters described in **A** to Astrocyte-like 2 (enriched in PERK KO) and S phase 2 populations (enriched in WT). **C**, Conditional dynamic differences for cluster 3 described in **A**. Top five genes are presented. **D**, Flow cytometric analysis of BNIP3 and PDK1 in healthy control (WT) and PERK-KO (KO) neurospheres. Left: Representative result, Right: Mean fluorescence intensity (MFI) (n =3). **E**, Relative levels of PFKFB4 in PFKFB4-knockout neurospheres. **F**, Flow cytometric analysis of BNIP3 and PDK1 in PFKFB4-knockout neurospheres. NTC = treated with negative control RNAi. Left: Representative result, Right: Mean fluorescence intensity (MFI) (n =3). Data are presented as the mean ± SD. ^*\**^*P <* 0.05, ^*\*\**^*P <* 0.01, ^*\*\*\**^*P <* 0.001.

### 3.5 Multiomics analysis of human hematopoiesis

The simultaneous profiling of transcriptomes and chromatin accessibility has been a standard technology to elucidate epigenetic regulations of transcriptional processes [6, 36]. Several studies have shown that chromatin accessibility changes prior to changes in gene expression [4, 36]. While the chromatin accessibility also describes the internal cell state as well as transcriptional profile, the versatility of ExDyn allow us to elucidate the regulatory effects of epigenetic states on transcriptome dynamics from the single-cell mutiomic data. Here, we calculated transcriptome factor (TF) activity [42] from chromatin accessibility of single-cell multiomic observations of hematopoiesis in human cord blood [38] and conditioned single-cell transcriptome dynamics by them. First, we confirmed that the latent cell states recapitulated the expected cell populations, such as hematopoietic stem cells, T cells, NK cells, B cells, and myeloid cells, and that the estimated cell state dynamics were directed from the stem cell population toward each cell lineage (**Fig. 5-A**). We explored the relationship between chromatin accessibility and the estimated cell state dynamics. To determine the magnitude of the dynamics changes induced by epigenetic differences, we calculated the Frobenius norm of the Jacobian matrix of cell state dynamics differentiated by TF activity for each cell. The influence of chromatin activity on the cell state dynamics of hematopoietic stem cells and progenitor cells was greater than that of most mature cells, such as T cells, B cells, and myeloid cells (**Fig. 4 in Supplementary**) .

**Figure 5:**
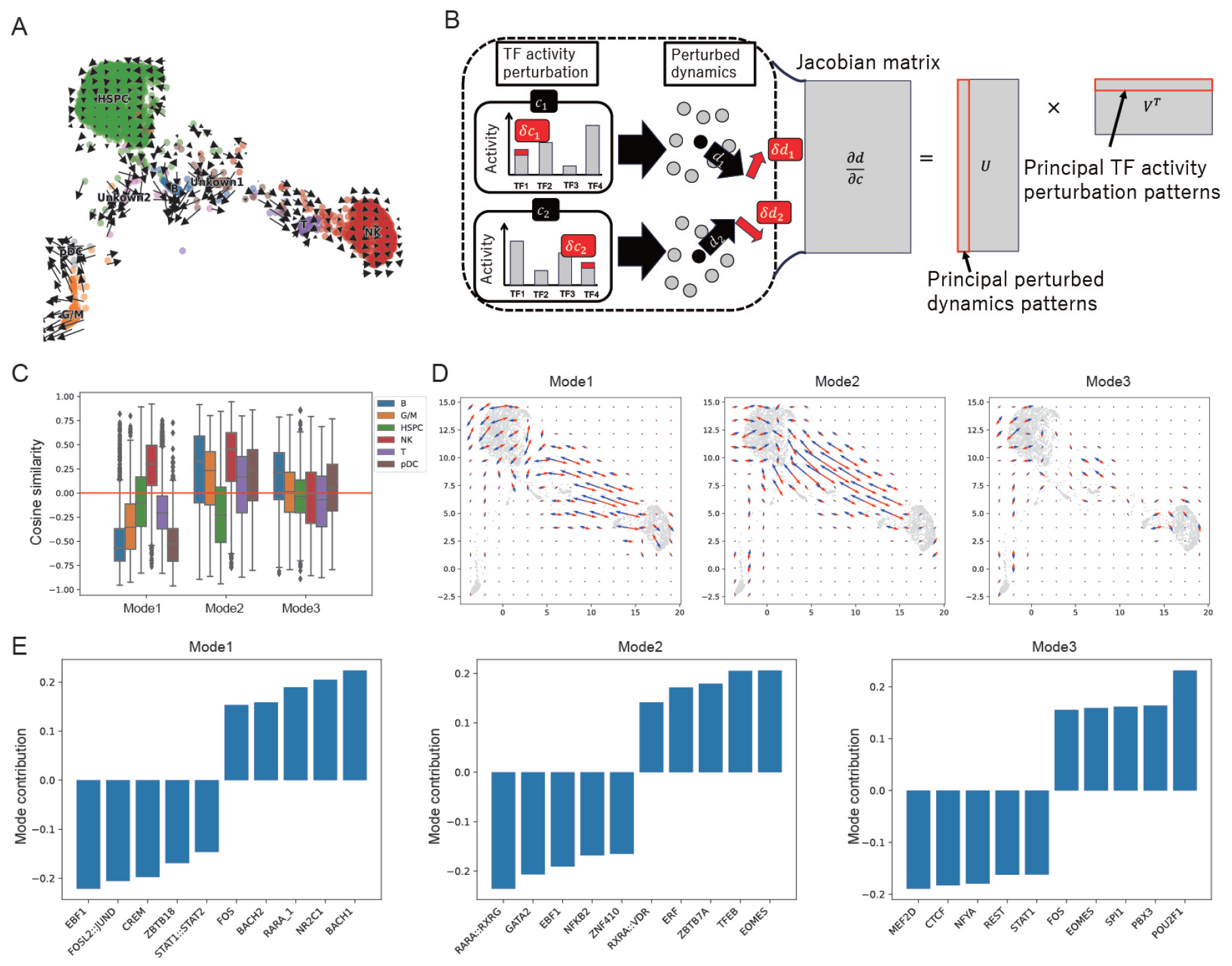
Chromatin-dependent cell state dynamics during human hematopoiesis. **A** 2D visualization of cell state dynamics in UMAP representation based on original TF activity. **B**, Schematic overview of perturbation analysis of cell state dynamics by TF activity. It extracts principal perturbation patterns of TF activity and contribution of each TF to each principal perturbation patterns. **C**, Cosine similarity between the cell state difference to each cell type and the top 3 modes of principal dynamic changes. **D**, 2D visualization of top 3 modes of principal dynamics changes. Red and blue arrows indicate the dynamics changes induced by positive and negative principal perturbations respectively. **E**, TF activity changes of top and bottom five contributing TFs for top 3 modes of principal dynamic changes.

We investigated the complex relationships between the activity of 125 TFs and transitions between cell states based on the SVD of the Jacobian matrix of gene expression change *v* differentiated by TF activity *c*, which gives 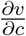 Briefly, the *n*-top mode of the SVD is based on the extrinsic factor change *δc*_*n*_, which maximizes the norm of the dynamic changes under the orthogonal condition to those of 1 *∼ n−*1-top modes and the corresponding change in cell state dynamics *δv*_*n*_ (see **Fig. 5-B** and ‘Singular value decomposition of dynamics Jacobian matrix’ in Methods).

We analyzed the direction of cell differentiation, which was enhanced when chromatin accessibility was changed along the direction of *c*_*n*_(*n* = 1, …, 5). We found that a positive change along *c*_2_ and *c*_3_ enhanced differentiation toward NK and B cell lineages, respectively, whereas a negative change along *c*_3_ enhanced differentiation toward the B and plasmacytoid dendritic cell lineages (**Fig. 5-C,D**). This suggests that different modes of changes in chromatin accessibility can induce differentiation into various cell lineages. We explored the transcription factors associated with chromatin changes in each mode. The transcription factor with the top contribution to the negative change along *c*_1_ was EBF1, whereas that to a positive change along *c*_2_ was EOMES1 (**Fig. 5-E**). Interestingly, EBF1 is a critical factor in B cell development [31], whereas EOMES1 regulates the development and maturation of NK cells [44]. The consistency between the enhanced differentiation direction and transcription factors with the highest contribution to the top modes of changes in chromatin accessibility demonstrated that ExDyn succeeded in capturing the complex relationships between cell state dynamics and chromatin accessibility.

### 3.6 ExDyn identifies fibroblasts inducing cancer cell invasion

Finally, we applied ExDyn to cancer cell populations in a combined single-cell and spatial transcriptome dataset of human squamous cell carcinoma [24] to investigate the effect of cell–cell communication within the tumor microenvironment on cell state transitions. We conditioned the cancer cell state dynamics by their colocalization strength with other cells, which was estimated using DeepCOLOR [27]. This conditioning assumes that the cell state transition is affected by the state of the neighboring cells. First, we confirmed that the cell state transitions were directed from cycling tumor keratinocytes toward basal, differentiated, and tumor-specific keratinocytes (**Fig. 6-A**), which is consistent with the fact that differentiated keratinocytes originate from proliferative keratinocytes [13]. We investigated the dependency of cell state dynamics on colocalization relationships using the SVD of the Jacobian matrix of cell state dynamics differentiated by colocalization profiles (see the previous section or ‘Singular value decomposition of dynamics Jacobian matrix’ in Methods). We found that the second mode of the colocalization change biased the cell state dynamics toward tumor-specific keratinocytes (TSKs), which were defined as malignant cancer cells with invasive features in the original publication [24], whereas the first mode enhanced differentiation toward differentiated keratinocytes (**Fig. 6-B,C**). To identify the colocalized cells that enhance the cell state transition to TSKs, we analyzed the contribution of each to condition changes in the second mode. We found that half of the top 10% of contributing cells were fibroblasts, the subpopulations of which are often associated with tumor malignancy [39] (**Fig. 6-D, E**).

**Figure 6:**
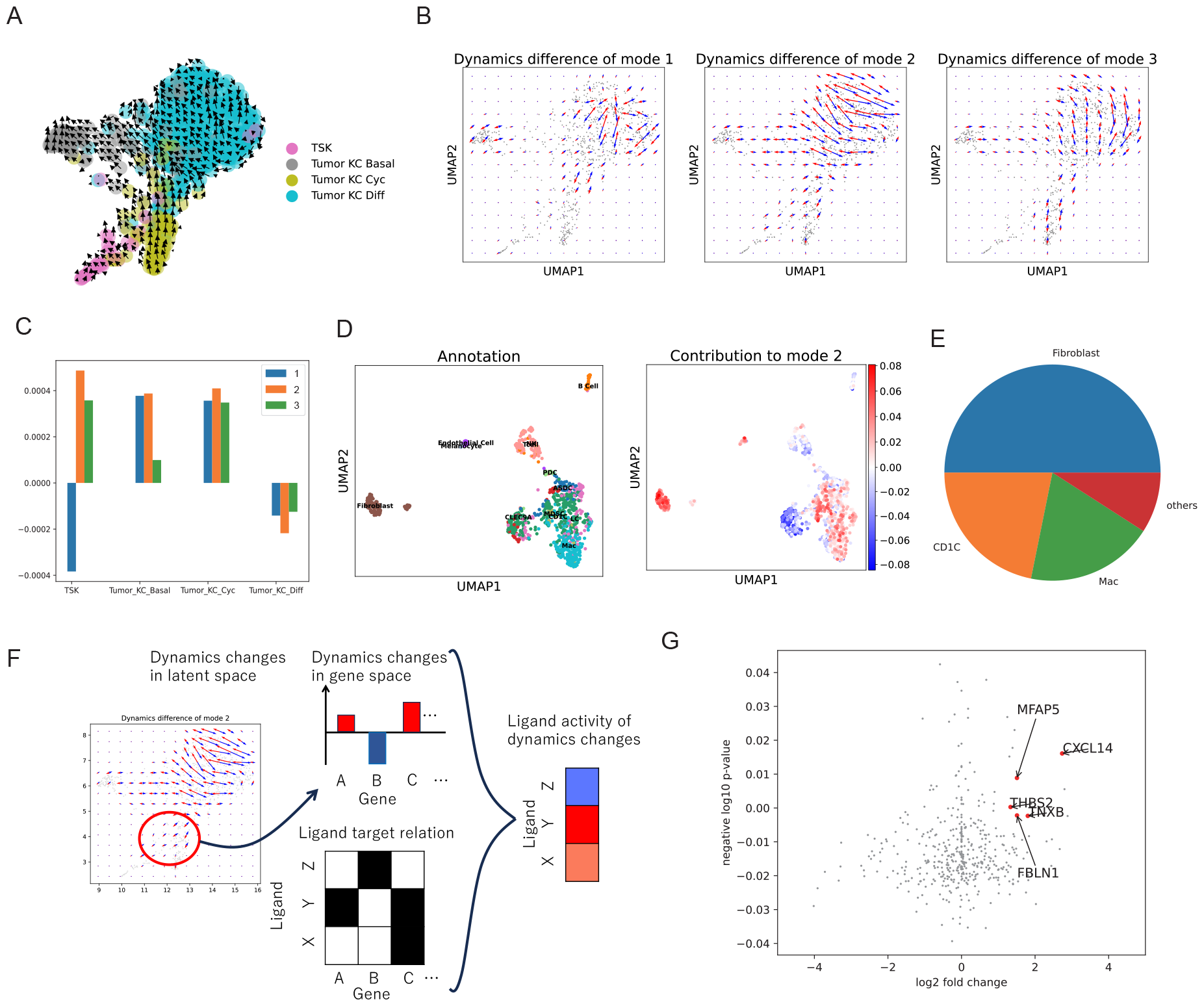
Environmental factors driving tumor heterogeneity in human skin squamous cell carcinoma. **A**, 2D visualization of cell state dynamics in UMAP representation based on original colocalization profiles. **B**, 2D visualization of top 3 modes of principal dynamics changes. Red and blue arrows indicate the dynamics changes induced by positive and negative principal perturbations respectively. **C**, Abundance changes in cancer subtypes induced by the top 3 modes of principal dynamic changes. **D**, Contribution to the mode 2 principal dynamics changes for each colocalized cells (right). The left panel represents cell type annotation which were derived from original publication. **E**, Proportions of cell types among the top-10% of cells contributing to the mode 2 principal dynamic differences. **F**, Schematic overview of the calculation of ligand activity for dynamics changes to TSK population. **G**, Ligand activity for mode 2 principal dynamic changes and log2 fold changes between fibroblasts with the top-10% contribution to mode 2 dynamic changes and the other fibroblasts. We highlighted the top-five most active ligands among the significantly upregulated ligands.

We investigated the molecular machinery regulating the cell state transition into TSKs induced by colocalization changes with fibroblasts based on the comparison between target genes of ligand signaling and genes with large dynamics changes *δv*_2_ induced by the mode 2 perturbation *δc*_2_. We summarized the dynamics changes in the targets for each ligand and conducted differential gene expression analysis of the ligands between colocalized fibroblasts and other cells (see **Fig. 6-F** and ‘Ligand activity for dynamics changes’ in the Methods). CXCL14, MFAP5, and THBS2 exhibited the top-3 largest positive changes in target gene dynamics among the significantly upregulated genes in this fibroblast population (adjusted p-value *<* 0.1 and log_2_ fold changes *>* 1) (**Fig. 6-G**). Indeed, MFAP5 is secreted by fibroblasts and induces invasive phenotypes in breast cancer cells [8]; CXCL14 and THBS2 have been reported to contribute to cancer invasion [30, 49]. These findings demonstrate the ability of ExDyn for identifying the molecular mechanisms underlying tumor cell heterogeneity dependent on colocalized cells.

### 3.7 Discussion

Existing computational methods for single-cell transcriptomes have focused on analyzing condition-specific populations [20,40] and the relationships between the transcriptome and extrinsic factors or covariates using co-embedding g [1, 18, 21, 35, 37]. However, it remains unclear how the relationships between extrinsic factors and the transcriptome arise. Conditioning the dynamics by extrinsic factors, the framework proposed in this study can elucidate the process by which specific cell populations are generated under varying extrinsic factors. This approach allowed us to directly capture the relationship between extrinsic factors and transcriptome dynamics and elucidate the mechanisms that conventional methods failed to reveal. Indeed, the factors identified with our approach are biologically plausible and can be experimentally validated, thus demonstrating the utility of ExDyn in revealing the dependency of gene regulatory mechanisms on specific extrinsic factors.

Splicing kinetics has enabled the data-driven estimation of RNA velocity and direction of cell state transitions [3, 28]. Although dynamic estimation without a presumptive differentiation direction can elucidate the generation process of specific cell populations, many existing methods rely on the smoothing of gene levels among similar cells, which complicates the analysis of cell dynamics between similar populations under varying extrinsic factors. We used a deep generative model to predict the unspliced transcriptome from extrinsic factors and the spliced transcriptome based on cell state dynamics. This approach effectively identified the relationships between extrinsic factors and the dynamics of each cell state and thus represents a new research avenue in cell state dynamics.

This study focused on cell state transition during a micro duration, which is an inherent nature of RNA velocity, and did not consider population shifts for a specific period of development or disease state. However, several studies have inferred transition, growth, and death across cell states using the time-course of population shifts [15, 23, 43, 52]. Hence, one possible extension of ExDyn is the inference of growth and death rates dependent on extrinsic factors, which can be realized by optimizing the time-course of the extrinsic factor-dependent prior distribution on the latent cell states from the observed time-course population shifts. We believe that ExDyn can be applied to develop a systematic approach for analyzing complex relationships between environmental factors and heterogeneous cell populations.

## 4 Methods

### 4.1 Deep generative model for extrinsic factor-dependent cell state dynamics

Here, we introduce our deep generative model for inferring extrinsic factor-dependent cell state dynamics. First, we introduce the generative process of spliced and unspliced transcriptomes in a manner consistent with splicing kinetics and cell state dynamics. Next, we present an algorithm for the variational inference of extrinsic factor-dependent cell state dynamics and the cell state itself.

#### 4.1.1 Generative process of spliced and unspliced transcriptome

For the spliced transcriptome, we applied the same approach as existing deep generative models of the single-cell transcriptome based on the VAE [12, 14, 32, 34]. Based on scVI [32], we adopted a conditional VAE (cVAE) [26] to exclude batch effects from the latent cell state space. We assumed that the latent cell state of cell *n, z*_*n*_ *∈ R*^*D*^ followed a standard Gaussian distribution:

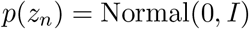

where *D* is the dimension of latent cell state space or Vamp prior [47]:

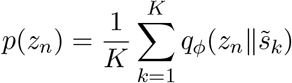

where *K* = 128 is the total number of mixture components, *s*_*k*_ is the *k*-th pseudo input of the spliced transcriptome (an optimizable parameter), and 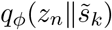 is the variational posterior distribution of cell states given the pseudo input (defined in ‘Variational inference of extrinsic factor-dependent dynamics’). We adopted a standard Gaussian distribution for the simulated and hematopoiesis datasets, and Vamp prior for the PERK KO and human skin squamous cell carcinoma datasets. We assumed that the size factor of cell *n, l*_*n*_, followed the lognormal distribution

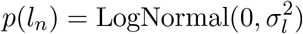

where 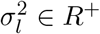 is an optimizable parameter. Spliced UMI counts of gene *g* in cell *c, s*_*c,g*_, follows a negative binomial distribution dependent on the latent cell state *z*_*n*_ and the experimental batches *b*_*n*_ *∈* {0, 1}^*B*^ where *B* is a total number of experimental batches and *b*_*n,k*_ = 1 if cell *n* belongs to experimental batch *k*:

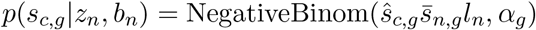

where 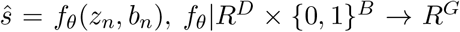 maps the latent cell states to the mean parameters of gene expression using a fully connected neural network (the architectures of the neural networks are presented in **Table. 1 in Supplementary** ), 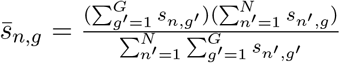 and *G* is the total number of genes. For the unspliced transcriptome, *u*_*n*_, we combined the latent cell state dynamics with the RNA velocity equation. Firstly, we assumed that the time evolution of cell states follows the simple diffusion process *dz* = *ρdW*, where *ρ* (set as 0.01 in this study) is the diffusion length over time. When noting the change in latent cell state during micro duration *δt* as *δtd*_*n*_, the instantaneous velocity *d*_*n*_ in the latent space follows the normal distribution

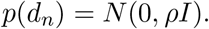

From the stochastic dynamics in the latent space, the instantaneous velocity in the gene expression space *v*_*n*_ can be calculated as follows:

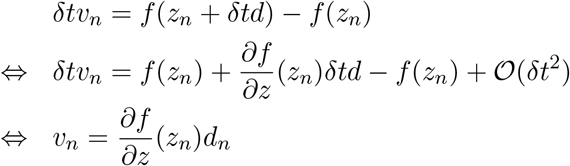

where the limit of *δt →* 0. Using the RNA velocity equation, we calculated the mean parameter of the unspliced transcriptome 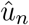 from the gene expression velocity *v*_*n*_ and the mean parameter of the spliced transcriptome 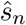 as follows:

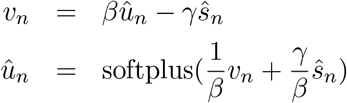

Because the mean parameter of the unspliced transcriptome is dependent on *z*_*n*_,*d*_*n*_,*l*_*n*_ and *b*_*n*_, we modeled the conditional distribution of the unspliced transcriptome as follows:

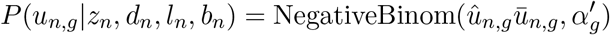

where *α*^′^ *∈ R*^+^ is an optimizable dispersion parameter, 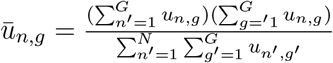 . In this model, the unspliced transcriptome is dependent on the stochastic variable for the dynamics in the latent space *d*. Hence, the posterior distribution of *d* mustis required to be consistent with the observed unspliced transcriptome, which is derived by from the variational inference described in the next section.

#### 4.1.2 Variational inference of extrinsic factor dependent dynamics

The generative process of unspliced and spliced transcriptomes includes two latent variables: latent cell state *z* and dynamics *d* . Here, we conducted variational inference in the similar manner with our previous work [38]. First, we assumed that the variational posterior distribution of *z* had a Gaussian distribution depending on the spliced transcriptome:

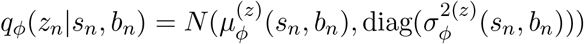

where 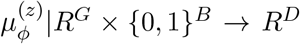 and 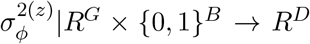 are neural networks for the mean and variance parameters of the variational distribution (the architectures of the neural networks are presented in **Table. 1 in Supplementary** ). We defined the variational posterior distribution for the latent cell state dynamics *d* as a Gaussian distribution depending on *z*_*n*_ and extrinsic factor *c*_*n*_ *∈ R*^*C*^ as follows:

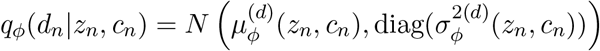

where *C* is the dimension of extrinsic factors, 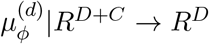 and 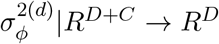 are neural networks for the mean and variance parameters of the variational distribution. Here, we noted that the mean and variance parameters of the latent cell state dynamics were functions of the latent cell state *z*_*n*_ and the corresponding extrinsic factors *c*_*n*_; hence, the relationships between the cell state dynamics and extrinsic factors can be effectively learned by the neural networks. We defined the variational posterior distribution for the library size*l*_*n*_ as a Log Normal distribution depending on *s*_*n*_ as follows:

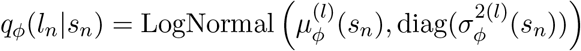

where 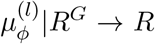 and 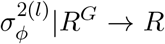 are neural networks for the mean and variance parameters of the variational distribution.

#### 4.1.3 Optimization of the generative model and variational posterior distribution

We optimized the generative model and variational posterior distribution by maximizing the evidence lower bound (ELBO), defined as follows:

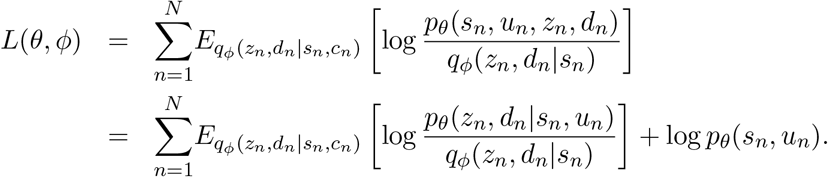

This objective function is equivalent to the summation of the marginal likelihood of the generative model and the Kullback (KL) divergence between the posterior distribution of the generative models and the variational posterior distribution. Hence, the ELBO maximization enhances the similarity between the variational and generative model posteriors, and increases the likelihood of the generative model. However, because the original ELBO is not analytically tractable, we transformed the original ELBO and conducted a Monte Carlo approximation in the same manner as the VAE as follows:

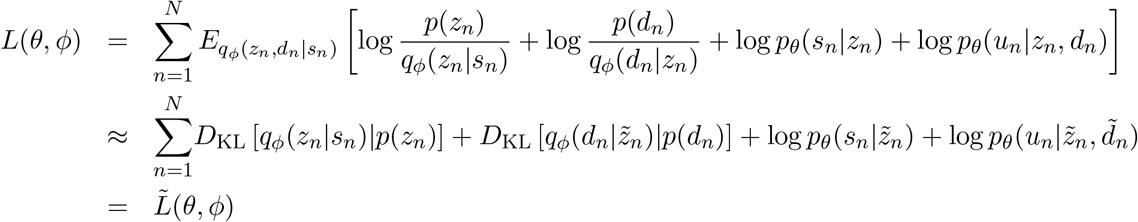

where 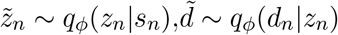 are differentiable by *ϕ*, since they are computed using a reparameterization [25]. We noted that the first and second terms were KL divergences between Gaussian distributions and therefore tractable for the standard Gaussian prior; we also adopted the combination of stochastic sampling of *z*_*n*_ and reparameterization to calculate the KL term in the Vamp prior.

We optimized *θ* and *ϕ* by minimizing the negative approximated ELBO 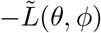 using the AdamW optimizer [33] in the PyTorch Library. We set the maximum number of epochs to 500, batch size to 128, and initial learning rate to 0.001. Using PytorchLightning library, we adopted early stopping with optional parameter ‘patience’ as 50 and ‘ReduceLROnPlateau’ as the learning rate scheduler with ‘patience’ as 20. We define the optimized parameters of *θ* and *ϕ* as *θ*^*\**^ and *ϕ*^*\**^ respectively.

### 4.2 Differential dynamics analysis for discrete conditions

We compared the cell state dynamics between two discrete conditions, *c*^(1)^ and *c*^(2)^. Using the optimized variational posterior distribution of cell state dynamics 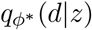, we first estimated the mean parameter of cell state dynamics and corresponding gene expression changes for each condition across cells as follows:

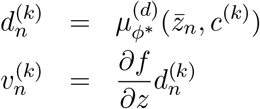

where *n ∈* {1, …, *N*} and *k ∈* {1, 2}. We analyzed the differences between conditions, which are denoted as 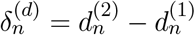 and 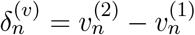.

### 4.3 Clustering analysis of populations with large dynamics difference between conditions

Condition difference can affect cell state dynamics for several subpopulations, and result in the different condition specific subpopulations. In order to analyze the generation process of the population difference, it is required to identify the population bifurcation points, the cell state dynamics of which are largely dependent on the conditions, and classified then in the unsupervised manner. Here, we firstly identified the cell populations with the top 10% largest L2-norm of the dynamics difference, ‖*δ*^(*v*)^‖, and conducted the clustering on the population using Leiden algorithm [48] implemented in Scanpy Python package [50]. We notated the cell members of *k*-th cluster as *C*_*k*_ *⊂* {1, … *N*} and the averaged difference of cell state dynamics and gene expression changes as 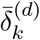 and 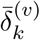. We note that we used 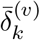 for the identification of genes with dynamics largely affected by the conditional difference.

### 4.4 Singular value decomposition of dynamics Jacobian matrix

Unlike batched experimental conditions, it remains difficult to analyze the dependency of cell state dynamics on continuous multivariate extrinsic factors, such as chromatin accessibility and cellular colocalization. We explored the conditional difference maximizing the dynamic difference, given that the norm of the conditional difference is 1. The dynamic difference *δ*^(*d*)^ induced by conditional difference *δ*^(*c*)^ was calculated as follows:

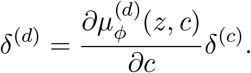

For the SVD of the Jacobian matrix 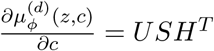, the norm of the dynamic difference can be formulated as follows:

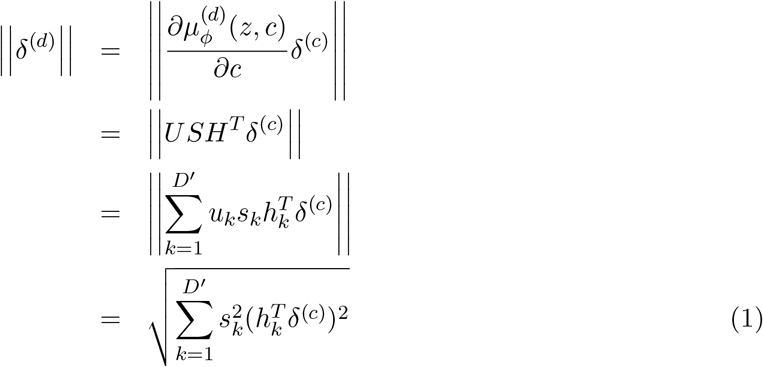

where *D*^*′*^ = min(*D, C*), *u*_*k*_ and *h*_*k*_ are the *k*-th column vectors of *U* and *V*, respectively, and*s*_*k*_ is the *k*-th singular value. Any conditional difference can be formulated using a linear combination of the column vectors of *H* and an orthogonal vector for them *h*^0^ as follows:

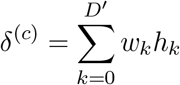

where the norm of the conditional difference is 1; hence, 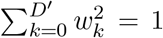. Substituting the linear combination into **Eq. 1**, the norm of the dynamic difference can be formulated as:

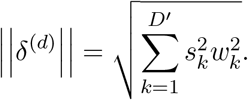

Owing to the weight constraint of the linear combination, the first column vector of *H, h*_1_, which is a linear combination with weight parameters *w*_1_ = 1, *w*_2_ = 0, …, *w*_*M*_ = 0, represents the conditional difference that maximizes the norm of the dynamic difference. Recursively, *m*-th column vector *h*_*m*_ is a vector maximizing the dynamic norm among the vectors orthogonal to *h*_1_, *h*_2_, …, *h*_*m−*1_ is the conditional difference that induces the *m*-th top dynamic difference *s*_*m*_*u*_*m*_, where *s*_*m*_ is *m*-th singular value and *u*_*m*_ is *m*th column vector of *U* .

### 4.5 Visualizing cell state dynamics in two-dimensional embeddings

In single-cell transcriptome analysis, it is often useful to visualize two-dimensional embeddings of cell states when elucidating the underlying biological processes. Existing methodologies project cell state dynamics into two-dimensional embeddings using the transition relationships between cells, calculated from the cell state dynamics. However, this transition-based approach is problematic because the embedded cell state dynamics are calculated by regarding the transition as the transition probability, while the transition quantify the similarity between RNA velocity and expression difference to target cells rather than the actual probability that the transition occurs; further, owing to the cell density in two-dimensional embeddings, similarities between dynamics and relative positional differences can bias this approach. Here, we propose an alternative approach for visualizing cell state dynamics using the chain rule of differentiation. First, we constructed a regression model of two-dimensional embedding based on the local weighted average of two-dimensional embedding *e ∈ R*^2^ as follows:

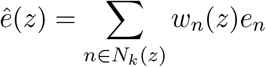

where *N*_*k*_(*z*) is *k*-nearest neighborhood of z*z* among *z*_*n*_(*n* = 1, …, *N* ) (*k* = 100 in this study), *e*_*n*_ is the two-dimensional embedding of the *n*-th cell and *w*_*n*_(*z*) is the weight of the *n*-th cell, calculated as follows:

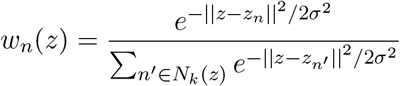

where *σ* is the bandwidth of the local weighted average. We set *σ* as the median of the distance between z and its *k*^*′*^-nearest neighbor ( 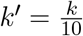 in this study). Using the chain rule of differentiation, we calculated the micro change in 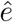 induced by the micro change in the cell state, *δ*_*z*_, as follows:

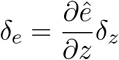

We visualized the cell state dynamics *d* and the dynamics difference *δ*^(*d*)^ using the change of 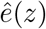 induced by them, 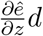 and 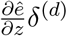.

### 4.6 Quantifying changes in abundance based on differential dynamics

We applied the two-dimensional embedding visualization to assess the changes in one-dimensional metrics on a cell-by-cell basis. Specifically, we quantified changes in the density of cluster *c* within the nearest neighborhoods due to the principal dynamics change *v*_*k*_ using variable *o*_*c*_, which represents the affiliation to cluster *c* and is normalized such that 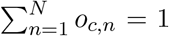, instead of *e*. We quantified the change in abundance due to dynamic changes in a particular cluster using the following score:

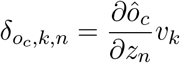

where

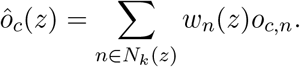

### 4.7 Preprocessing of scRNA-seq

For PERK-KO and WT neurospheres, we mapped the scRNA-seq reads to the human genome (GRCh38) using CellRanger. For all datasets, we quantified the unspliced and spliced transcriptomes using ‘kb-bustools’ with the corresponding reference genome and technology parameter (‘-x 10xv3’). We specified the optional parameter ‘workflow’ as ‘lammanno’. For all datasets, we conducted similar filtering procedures on cells and genes using the Python package ‘Scanpy’. First, we filtered cells and genes that did not satisfy the conditions listed in **Table. 2 in Supplementary** . We conducted total count normalization using ‘scanpy.pp.normalize total’, log2 transformation using ‘scanpy.pp.log1p’, and principal component analysis (PCA) using ‘scanpy.tl.pca’ for genes selected by ‘scanpy.pp.select_highly_variable_genes’ with ‘n_top_genes = 4000’. Using the PCA co-ordinates, we calculated k-nearest neighborhoods using ‘scanpy.pp.neighbors’ with ‘n_neighbors = 300’ and the moment of the spliced and unspliced transcriptomes using ‘scvelo.pp.moments’ in the Python package ‘scVelo’. We calculated the likelihood of an unspliced transcriptome using only genes with high Pearson’s correlation coefficients for spliced and unspliced transcriptome moments (*>* 0.6). This should improve the accuracy when estimating cell state dynamics.

### 4.8 Differential gene expression analysis

We conducted differential gene expression analysis using ‘sc.tl.rank_genes_groups’ in the Python package ‘Scanpy’ with ‘method = wilcoxon’. We identified differentially expressed genes based on adjusted p-value ¡0.1 and absolute log-fold change ¿1.

### 4.9 Preprocessing for chromatin accessibility

In the multiomics analysis of chromatin accessibility and the single-cell transcriptome in the human hematopoiesis dataset, we mapped both of the scRNA-seq and scATAC-seq reads to the human genome (GRCh38) using CellRanger-ARC. cell state dynamics were conditioned based on transcription factor activity calculated using the R package ‘chromVar’ [42] for improved interpretability. We used the Signac R package to calculate TF activity [46]. We fetched position frequency matrices using ‘getMatrixSet’ from JASPAR2020 [16] with the optional parameters ‘collection = CORE’, ‘tax_group = vertebrates’, and ‘all_versions = FALSE’ and generated motif information using ‘AddMotifs’ with the optional parameter ‘genome = BSgenome.Hsapiens.UCSC.hg38’. Finally, we calculated TF activity using ‘RunChromVAR’ with the optional parameter ‘genome = BSgenome.Hsapiens.UCSC.hg38’.

### 4.10 Colocalization estimation for cancer cells

In the skin squamous cell carcinoma dataset, we extracted cells derived from patient 6, since both of the single cell and spatial transcriptome analysis (Visium) was conducted for the patient. The Python package DeepCOLOR was used to estimate colocalization between cells, which was then used as a condition in the cell state dynamics estimation. After selecting genes with *>* 100 counts in both single-cell and spatial transcriptome observations, we estimated spatial distribution using ‘deepcolor.workflow.estimate_spatial_distribution’ and calculated the colocalization scores using ‘deepcolor.workflow.estimate_colocalization’. We extracted the cancer cell population using the annotation labels ‘Tumor_KC_Basal’, ‘Tumor_KC_Diff’, ‘Tumor_KC_Cyc’ and ‘TSK’ and excluded the colocalization with cancer cells themselves from the condition used in the dynamics inference because the colocalization scores were exceptionally high for nearly identical cells.

### 4.11 Ligand activity for dynamics changes

We quantified the activity of diverse ligand signals on dynamic changes to identify key environmental factors using ‘Nichenet’ [5] —a computational tool that calculates ligand activity based on signaling pathways and downstream gene regulation and provides a potential regulation score. We selected the top 1% of genes with this regulation score and averaged the dynamic changes in these genes to determine the activity of each ligand.

#### 4.11.1 Simulating differential cell state dynamics between discrete conditions

We used Sergio, a single-cell RNA-seq simulation based on a gene regulatory network (GRN) and a reaction kinetics model of transcription, splicing, and degradation to validate the estimation of different dynamics depending on extrinsic factors. Sergio simulates differentiation into specific cell types based on the transcription rates of the GRN and Master regulator. We used SERGIO/Demo/differentiation_input_GRN.txt, in the Sergio repository as the GRN, and the transcription rates for the master regulator were based on cell type 0 in SERGIO/Demo/differentiation_input_MRs.txt. For the source population, condition 1, and condition 2, we set the transcription rates of master regulator 79 to 1.0, 2.0, and 2.0; those of 18 to 1.0, 1.0, and 2.0; and those of 9 to 1.0, 2.0, and 1.0, respectively. The spliced and unspliced count data were generated based on the condition-specific transcription profile transitions associated with these transcription rate differences. To observe the cellular states during differentiation, we randomly sampled time points within a maximum time of 1000, instead of using the standard ‘getExpressions dynamics’ method implemented in Sergio. To benchmark the RNA velocity estimate, we calculated the changes in expression during the following 100 steps as the ground-truth RNA velocity.

#### 4.11.2 Compared RNA velocity methods

We compared the performance of ExDyn with that of the RNA velocity methods scVelo, VeloVI, and CellDancer using the default parameters for all methods. For scVelo, we used the ‘scv.tl.velocity’ function with ‘mode = dynamical’.

### 4.12 Human iPS cells (iPSCs) and neuronal induction

The PERK-deficient iPSC lines (PERK KO) were generated from healthy control iPSC line (WT, female) using CRISPR/Cas9 systems. The detail was described in our previous study [2]. They were differentiated into neurospheres as previously reported [2].

### 4.13 Single-cell RNA sequencing (scRNA-seq)

Neurospheres were dissociated using AccuMax (Thermo Fischer Scientific) and filtered to remove debris. Cells with viability ¿80% were used further analysis. Cell suspensions which target 10,000 cells were loaded into 10x chromium microfluidic devices to produced barcoded single-cell nanodroplet emulsions. After sample generation, barcoded-emulsions were broken, amplified, and libraries pre-pared according to manufacture instructions (Chromium Next GEM single Cell 3’ Reagent Kits v3.1 (Dual Index), 10x Genomics). Evaluation of cDNA was examined on a Bioanalyzer Systems (Agilent). Libraries were subsequently subjected to next generation sequencing (NovaSeq6000, illumina).

### 4.14 Knockdown of PFKFB4

PFKFB4 small interfering RNAs (siRNA) and negative control (NTC) siRNA were purchased from Sigma-Aldrich as follows: PFKFB4 siRNA#1: SASI_Hs01_00127438, PFKFB4 siRNA#2: SASI_Hs01_00127439, PFKFB4 siRNA#3 SASI_Hs02_00337894, and NTC siRNA SIC-001-10. The 10nM siRNAs were transfected into neurospheres using Lipofectamine RNAiMax (Thermo Fisher Scientific). After 96 hours of transfection, the cells were collected for further experiments. To evaluate the knockdown efficiency, the expression level of PEKFB4 mRNA was examined. Total RNA was extracted using the RNeasy Plus Mini Kit (QIAGEN). Reverse transcription was performed using a High-Capacity cDNA Transcription Kit (Applied Biosystems). qPCR-based gene expression analysis was conducted using THUNDERBIRD Next SYBR qPCR mix (TOYOBO) and QuantStudio5 (Applied Biosystems). The primers used were PFKFB4:*fw* TTTTTCTCCCCGACAATGAAGAG, *rv* CACACAGATGGACTCGACAAA and RPS18: *fw* GCGGCGGAAAATAGCCTTTG, *rv* GATCACACGTTCCACCTCATC.

### 4.15 Flow cytometry

Neurospheres were dissociated into single cells using AccuMax and then fixed in 4 % paraformaldehyde for 15 min, followed by treatment with ice-cold 90% methanol. After blocked in phosphate-buffered saline (PBS) containing 0.5 % BSA, cells were incubated with the indicated primary antibodies 1 hour on ice. After washing with PBS, immunolabeled cells were incubated with appropriate fluorophore-tagged secondary antibodies for 1 hour on ice. The cells were analyzed using BD FACSCantoII (BD). The primary antibodies used were anti-BNIP3 (ab10433, abcam) and anti-PDK1 (ab202468, abcam).

## Supporting information

Supplementary Information for ``Inferring extrinsic factor-dependent single-cell transcriptome dynamics using a deep generative model''

## 5 Data availability

scRNA-seq data of PERK KO and WT iPS derived neurosphere will be deposited in Gene Expression Omnibus. Combined spatial and single-cell transcriptome data of human SCC dataset is available in Gene Expression Omnibus (accession id: GSE144240). Single cell multiome data of human hematopoiesis data set is available in Japanese Genotype-phenotype Archive (accession id: JGAS000528). Codes for our analysis, including ExDyn, are available at https://www.github.com/kojikoji/exdyn.

## 6 Author contributions

YK designed the study and performed software development and data analysis. YA, KN, YM and HO conducted and supervised experiments. KO and YS supervised experiments. HH, SH, and MI conducted software development and data analysis. TS and NO designed and supervised the study.

## 7 Acknowledgement

This research was funded by multiple sources. The Grants-in-Aid for Scientific Research (B) (grant no. 20H04281), Grants-in-Aid for Scientific Research on Innovative Areas on Information Physics of Living Matters (grant no. 22H04839), Grant-in-Aid for Transformative Research Areas (platforms for Advanced Technologies and Research Resources) (grant no. 22H04925), Grant-in-Aid for Transformative Research Areas (A) (grant no. 23H04938), and Grant-in-Aid for Research Activity Start-up (grant no. 20K22839) were provided by the Japan Society for the Promotion of Science (JSPS). Additional support was received from RADDAR-J (grant no. JP22ek0109488), the Project for P-PROMOTE (grant nos. JP22ama221215 and JP22ama221501), Brain/MINDS Health and Diseases (grant no. JP22wm0425007), the Interdisciplinary Cutting-edge Research (grant no. JP23wm0325068), and the Advanced Genome Research and Bioinformatics Study to Facilitate Medical Innovation (GRIFIN) (grant no. JP23tm0424226) from the Japan Agency for Medical Research and Development (AMED). The Moonshot Moonshot R&D program (grant no. JPMJMS2025) and ACT-X program (grant no. JPMJAX20AB) also contributed, through the Japan Science and Technology Agency (JST). Further support came from the Medical Research Center Initiative for High Depth Omics and Multilayered Stress Diseases at Tokyo Medical and Dental University. Supercomputing resources were provided by the Shirokane supercomputer at the Human Genome Center of the University of Tokyo, the TSUBAME3.0 supercomputer at the Tokyo Institute of Technology, and the AI Bridging Cloud Infrastructure (ABCI) at the National Institute of Advanced Industrial Science and Technology (AIST).

## 8 Competing interests

The authors declare no competing interests.

## Notes

### Competing Interest Statement

The authors have declared no competing interest.

