## Supplementary Information for ``Inferring extrinsic factor-dependent single-cell transcriptome dynamics using a deep generative model'' for "Inferring extrinsic factor-dependent single-cell transcriptome dynamics using a deep generative model"

Table 1: ExDyn architecture

| Name | Operation | Dimension | Normalization | Activation | Input |
|---|---|---|---|---|---|
| **Inputs** | | | | | |
| SPLICED | - | #Genes | - | - | - |
| COV | - | #Covariates | - | - | - |
| BATCH | - | #Batches | - | - | - |
| **Encoder** $z$ | | | | | |
| $h_1^{(z)}$ | FC | 128 | Layer | ReLU | SPLICED, BATCH |
| $h_2^{(z)}$ | FC | 128 | Layer | ReLU | $h_1^{(z)}$ |
| $\mu_\phi^{(z)}(x)$ | FC | 10 | No | - | $h_2^{(z)}$ |
| $\sigma_\phi^{2(z)}(x)$ | FC | 10 | No | Softplus | $h_2^{(z)}$ |
| $z$ | Sample from Normal | 10 | - | - | $\mu_\phi^{(z)}(s)$, $\sigma_\phi^{2(z)}(s)$ |
| **Encoder** $d$ | | | | | |
| $h_1^{(d)}$ | FC | 128 | Layer | ReLU | SPLICED, COV |
| $h_2^{(d)}$ | FC | 128 | Layer | ReLU | $h_1^{(d)}$ |
| $\mu_\phi^{(d)}(x)$ | FC | 10 | No | - | $h_2^{(d)}$ |
| $\sigma_\phi^{2(d)}(x)$ | FC | 10 | No | Softplus | $h_2^{(d)}$ |
| $z$ | Sample from Normal | 10 | - | - | $\mu_\phi^{(d)}(z)$, $\sigma_\phi^{2(d)}(z)$ |
| **Decoder s** | | | | | |
| $h_1^{(s)}$ | FC | 128 | Layer | ReLU | z, BATCH |
| $h_2^{(s)}$ | FC | 128 | Layer | ReLU | $h_1^{(s)}$ |
| $f_\theta(z)$ | FC | #Genes | No | Softplus | $h_2^{(s)}$ |

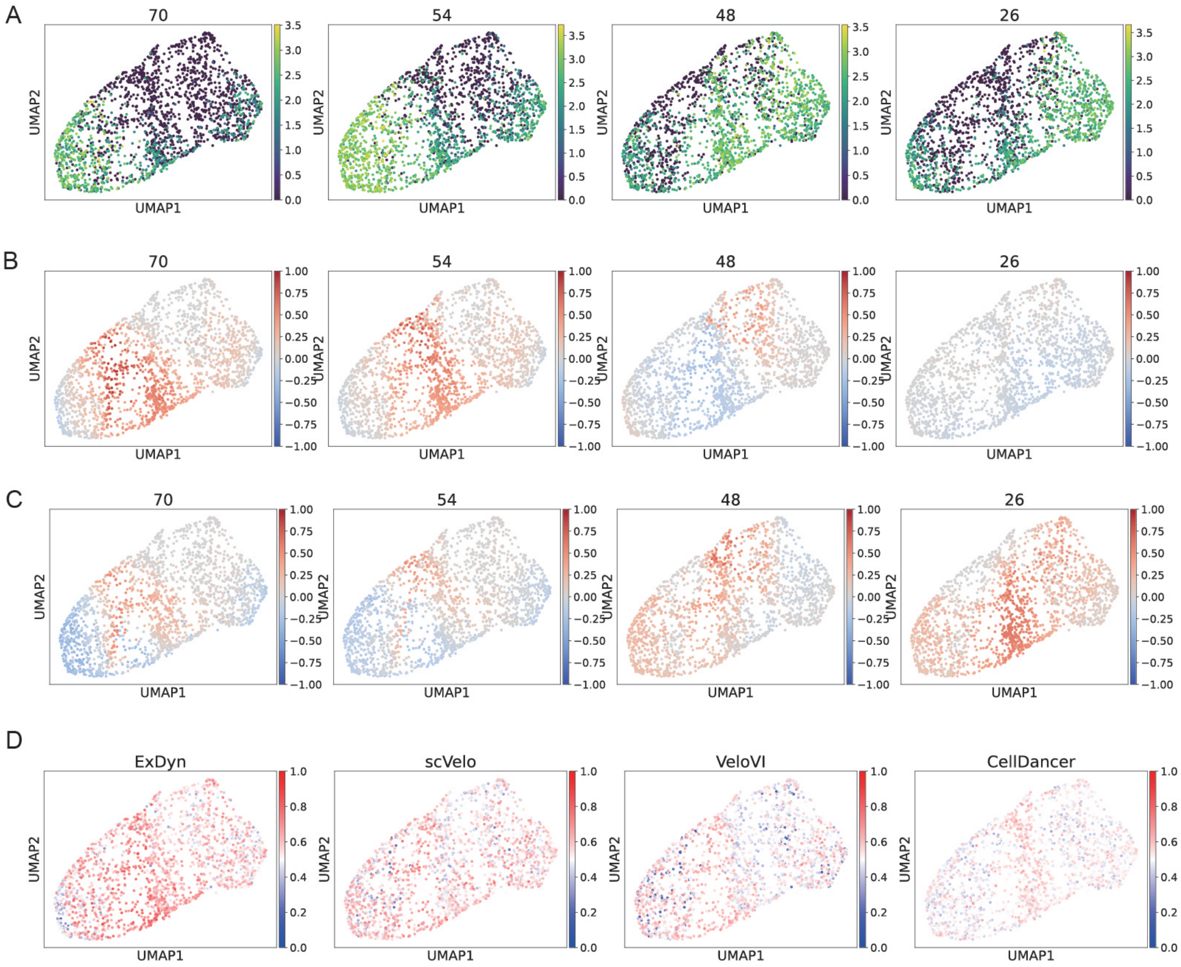


Figure 1: **Condition-wise gene expression velocity in simulated dataset.** We displayed gene expression (**A**) and velocity in condition 1 (**B**) and condition 2 (**C**) for genes with top 2 up and down regulated genes in condition 1 compared to condition 1. **I**, Cell-wise accuracy of RNA velocity estimated using ExDyn and other RNA velocity methods (scVelo, VeloVI, and CellDancer). The accuracies were defined as the proportion of genes with consistent signatures between estimated and ground-truth RNA velocities for each cell.

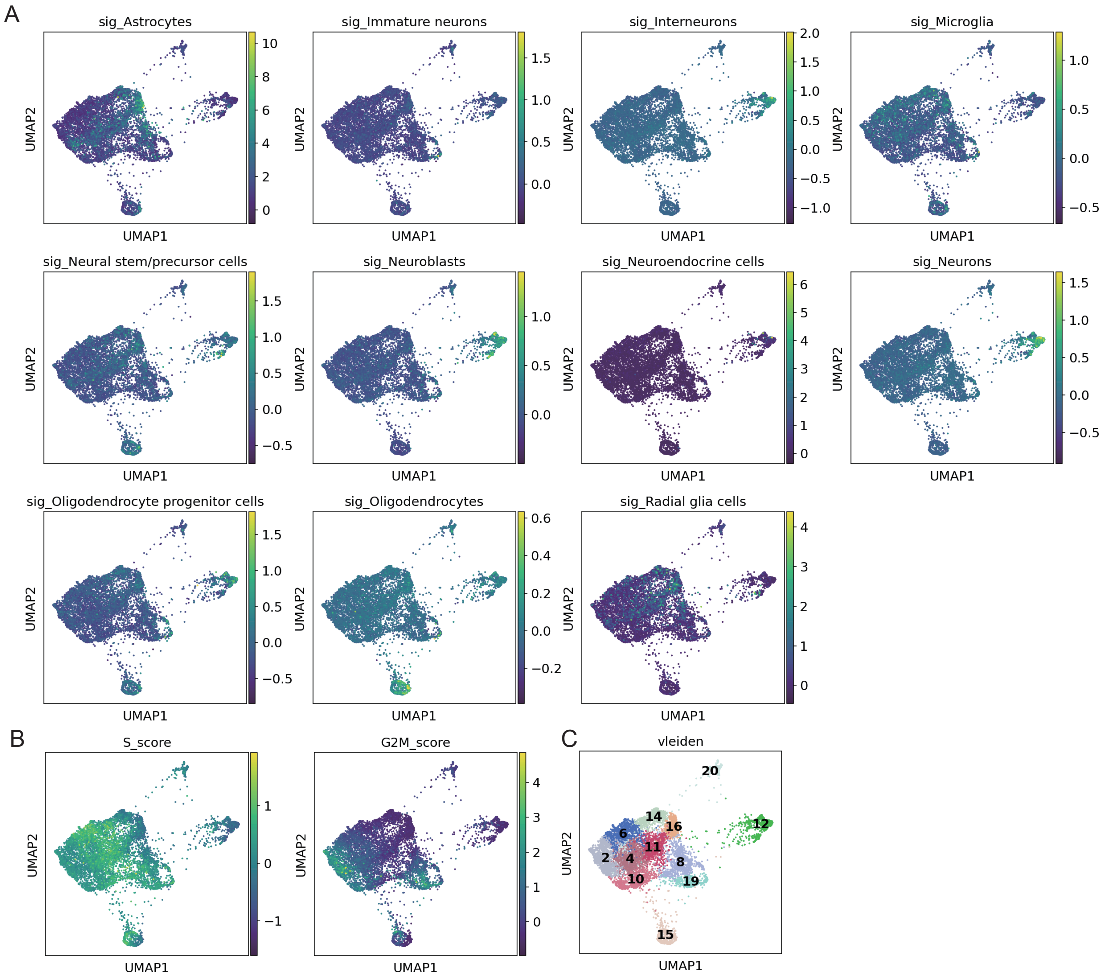


Figure 2: **Annotation on PERK KO Neurosphere dataset. A** Signature scores of marker genes for cell types belonging to brain tissues. **B**, Signature scores for S phase and G2M phase specific genes. **C**, Clusters derived by Leiden algorithm.

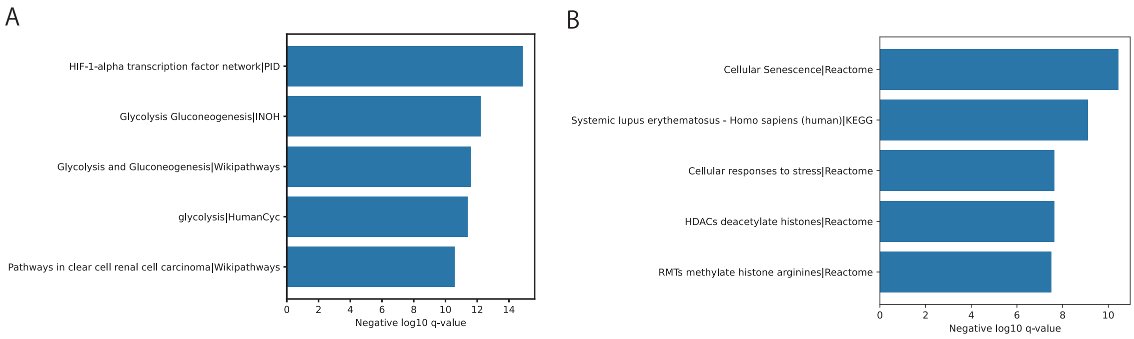


Figure 3: **Pathway enrichment analysis on condition specific population** Pathway enrichment analysis on up regulated genes (log2 fold changes $> 1$ and adjusted p values $< 0.05$) for Astrocyte like 2 (**A**) and S phase 1 **B**.

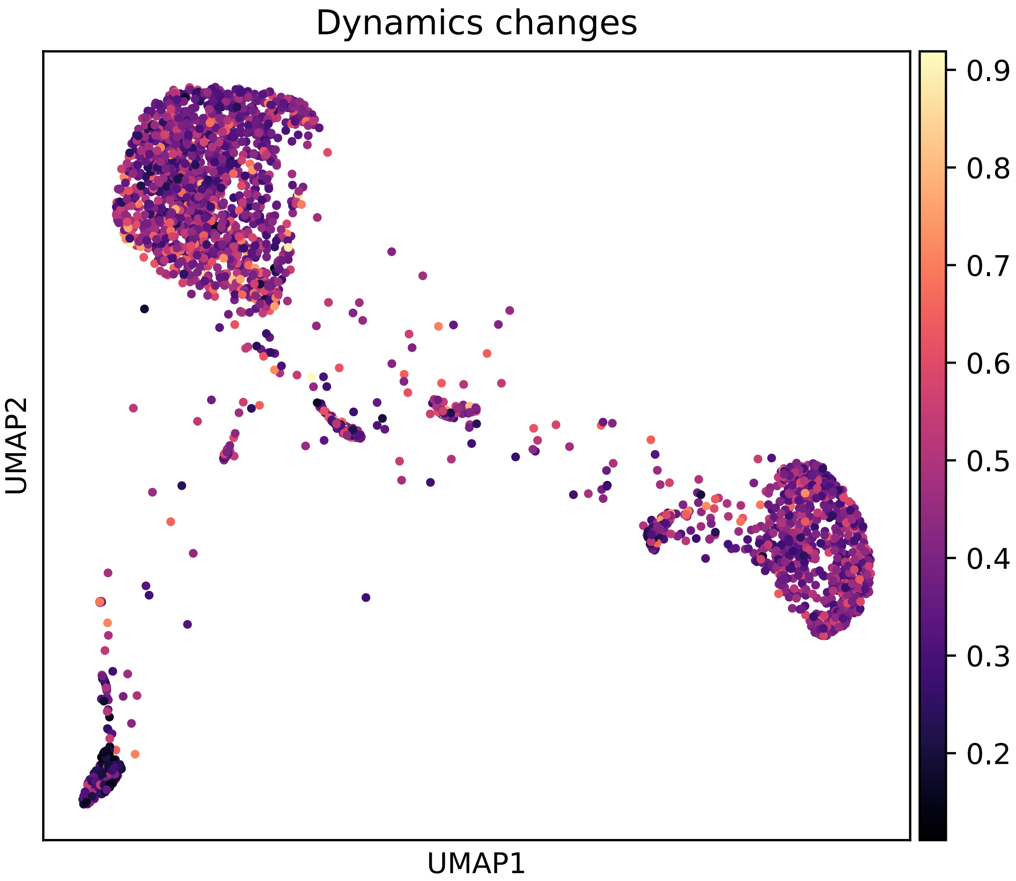


Figure 4: **Perturbation strength on cell state dynamics by chromatin activity** Magnitude of dynamic changes induced by perturbations in TF activity, which is defined by the Frobenius norm of the Jacobian matrix of cell state dynamics differentiated by TF activities.
